## supplemental information for "Center-surround inhibition in expectation and its underlying computational and artificial neural network models"

1. **Figure 2 – Figure supplement 1 | Accuracies of auditory tone report in the profile experiment.**
2. **Figure 3 – Figure supplement 1 | The calculated parameters of the population responses on OD and SFD tasks.**
3. **Figure 3 – Figure supplement 2 | Results of Combined computational modeling.**
4. **Figure 5 – Figure supplement 1** **| Accuracies of auditory tone report in orientation adjustment experiments.**
5. **Figure 6 – Figure supplement 1 | Adjusted errors of orientation adjustment experiments and their parameter estimates with the three-component mixture model.**
6. **Figure 7 – Figure supplement 1 | Accuracies of auditory tone report in orientation discrimination experiments.**

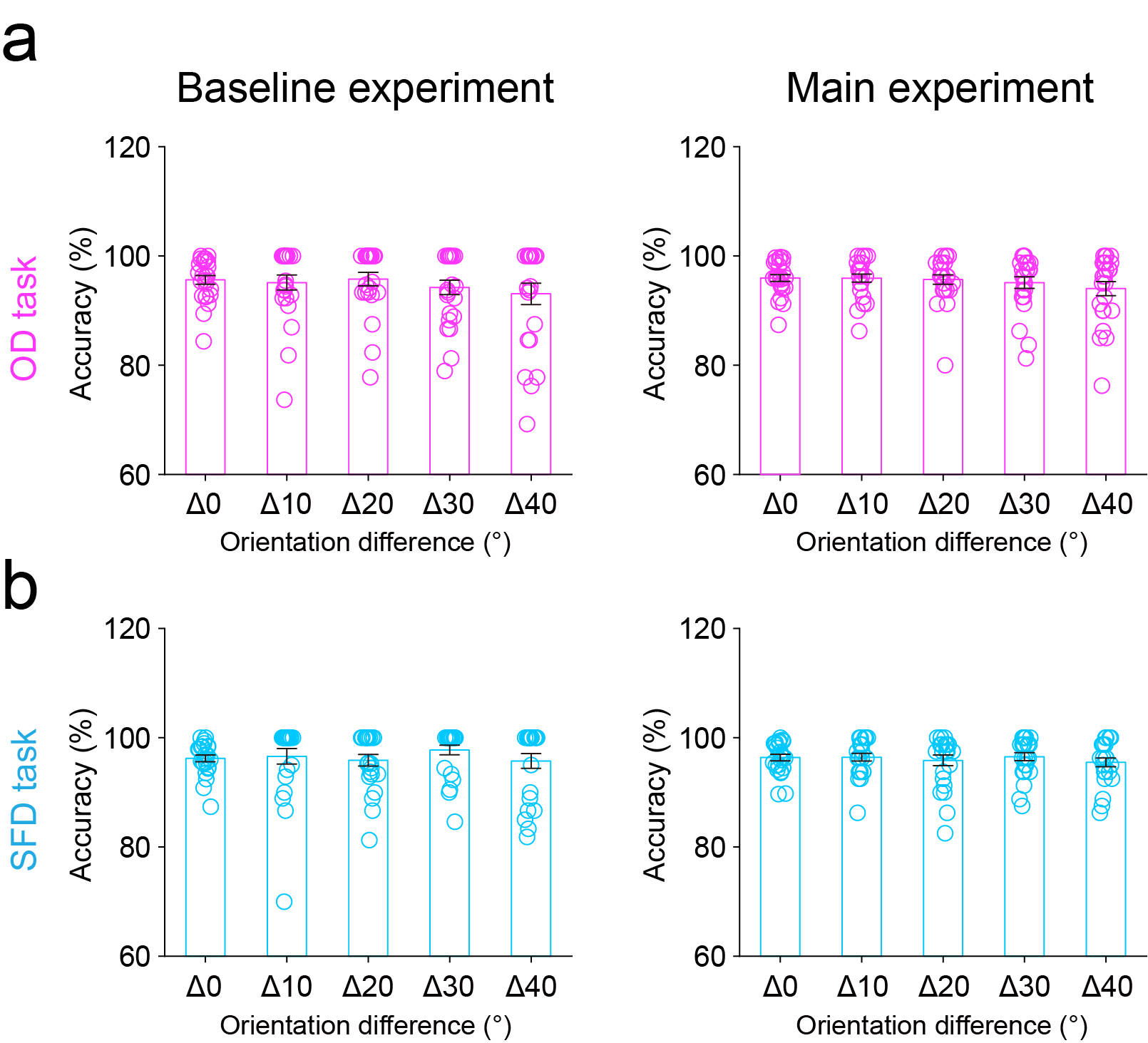

**Figure 2 – Figure supplement 1 | Accuracies of auditory tone report in the profile experiment.** Mean accuracies of auditory tone reports in OD (**a**) and SFD (**b**) tasks during the baseline (left) and main (right) experiments, plotted across different distances in orientation space (Δ0° - Δ40°). In both the baseline and main experiments, a one-way repeated-measures ANOVA with the distance (Δ0°-Δ40°) as within-participants factor showed that, the main effect of distance was not significant for either OD (baseline: F(4,92) = 1.281, p = 0.283, 𝜂_p_^2^ = 0.053; main: F(4,92) = 2.412, p = 0.082, 𝜂_p_^2^ = 0.095) or SFD (baseline: F(4,92) = 0.631, p = 0.594, 𝜂_p_^2^ = 0.027; main: F(4,92) = 0.963, p = 0.414, 𝜂_p_^2^ = 0.040) tasks. Error bars indicate 1 SEM calculated across participants.

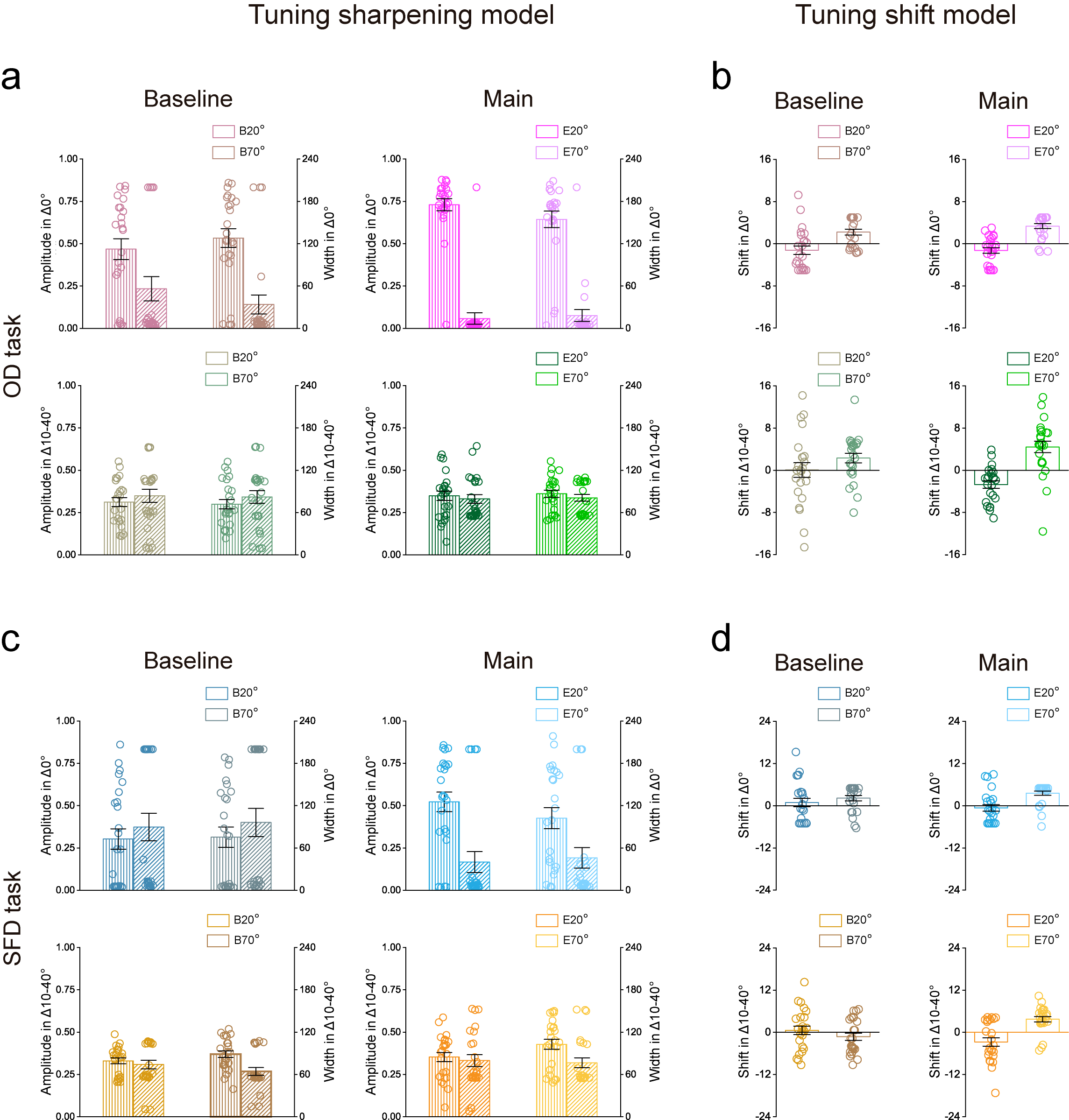

**Figure 3 – Figure supplement 1 | The calculated parameters of the population responses on OD and SFD tasks**. **a,** Amplitude *A* (vertical stripes) and Width *σ* (diagonal stripes) of the Tuning sharpening model for the baseline (left) and main (right) experiments in the Δ0° (top) and Δ10°-Δ40° (bottom) conditions. Error bars indicate 1 SEM calculated across participants. **b,** Location *x0* of the Tuning shift model for the baseline (left) and main (right) experiments in the Δ0° (top) and Δ10°-Δ40° (bottom) conditions*.* Note that the data were presented as the bias between *x0* and their hypothesized channel location, i.e., 20°, 30°, 40°, 50°, 60°, and 70°. Error bars indicate 1 SEM calculated across participants. **c** and **d,** Computational modeling for the SFD task, see caption for (**a** and **b**) for a description of each type of graph

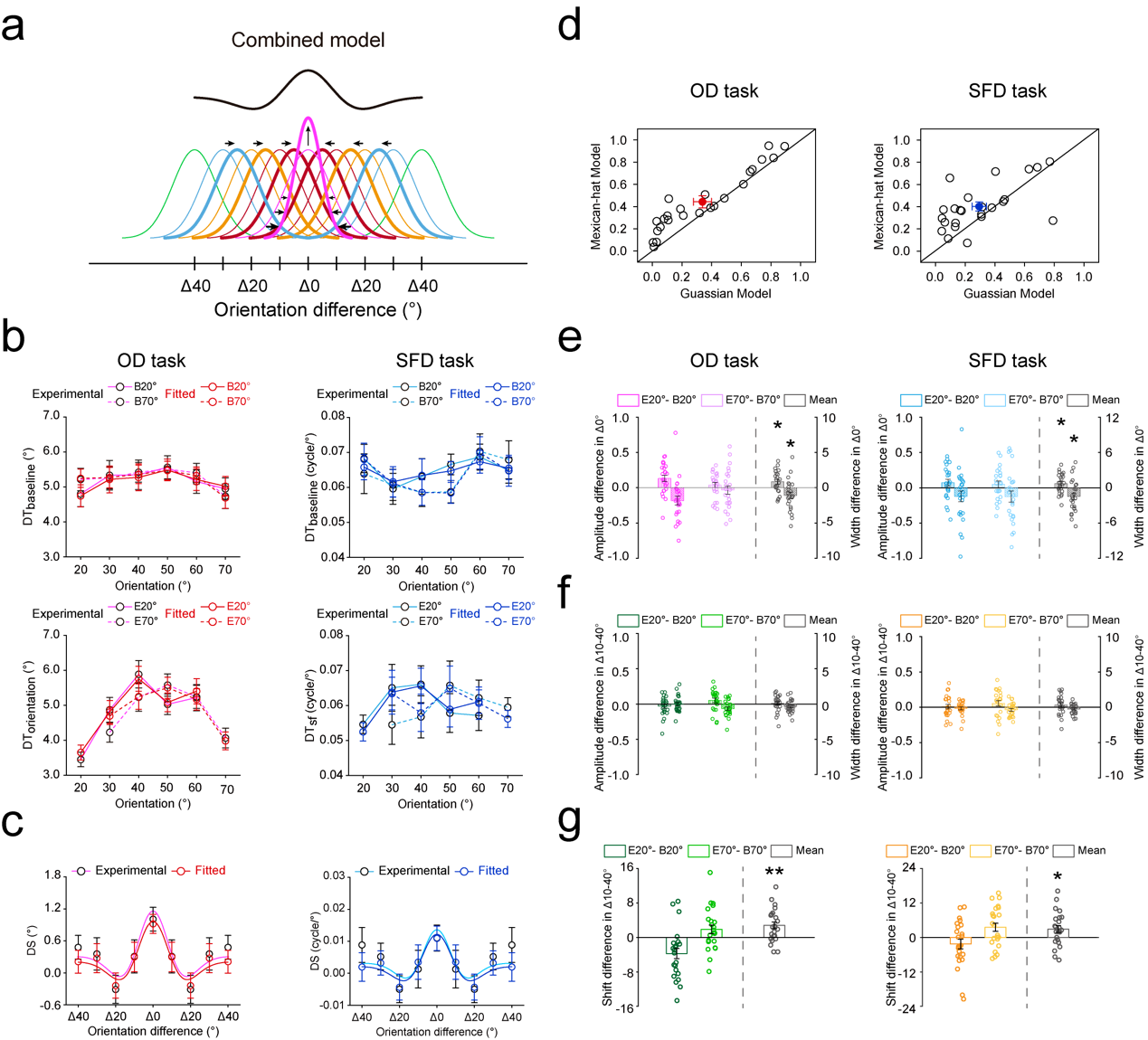
 **Figure 3 – Figure supplement 2 | Results of combined computational modeling. a,** Illustration of the Combined model. The Combined model incorporates sharpening of the expected orientation channel (center channel) together with shifting of the unexpected orientation channels (surround channels) from unexpected toward the expected orientation, resulting in a center–surround population response profile (black curve). **b,** The fitted discrimination thresholds on OD (left) and SFD (right) tasks in the baseline (top) and main (bottom) experiments. **c,** The averaged DSs using Combined model on OD (left) and SFD (right) tasks. **d,** R^2^ of the best fitting Gaussian and Mexican-hat functions for individual participants based on the fitted DSs on OD (left) and SFD (right) tasks. Open symbols represent the data from each participant and filled colored dots represented the mean across participants. The amplitude *A* (vertical stripes) and width *σ* (diagonal stripes) differences between the baseline and main experiments in Δ0° (**e**) and Δ10°-Δ40° (**f**) conditions, on OD (left) and SFD (right) tasks. The location *x0* differences between the baseline and main experiments Δ10°-Δ40° (**g**) conditions, on OD (left) and SFD (right) tasks. For the expected orientation (Δ0°) , results showed that the amplitude change was significantly higher than zero on both OD (t(23) = 2.582, p = 0.017, Cohen’s d = 0.527) and SFD (t(23) = 2.078, p = 0.049, Cohen’s d = 0.424) tasks (**e**, vertical stripes); the width change was significantly lower than zero on both OD (t(23) = -2.438, p = 0.023, Cohen’s d = 0.498) and SFD (t(23) = -2.578, p = 0.017, Cohen’s d = 0.526) tasks (**e**, diagonal stripes). For unexpected orientations (Δ10°-Δ40°), however, the amplitude and width changes were not significant with zero on either OD (amplitude change: t(23) = 0.443, p = 0.662, Cohen’s d = 0.091; width change: t(23) = -1.819, p = 0.082, Cohen’s d = 0.371) or SFD (amplitude change: t(23) = 1.130, p = 0.270, Cohen’s d = 0.231; width change: t(23) = -1.710, p = 0.101, Cohen’s d = 0.349) tasks (**f**). In the meantime, the location shift was significantly different than zero for unexpected orientations (Δ10°-Δ40°, OD task: t(23) = 3.611, p = 0.001, Cohen’s d = 0.737; SFD task: t(23) = 2.418, p = 0.024, Cohen’s d = 0.493 (**g**). Open symbols represent the data from each participant and error bars indicate 1 SEM calculated across participants. B20°: Baseline 20°; B70°: Baseline 70°; E20°: Expect 20°; E70°: Expect 70° (*p < 0.05; **p < 0.005; ***p < 0.001).

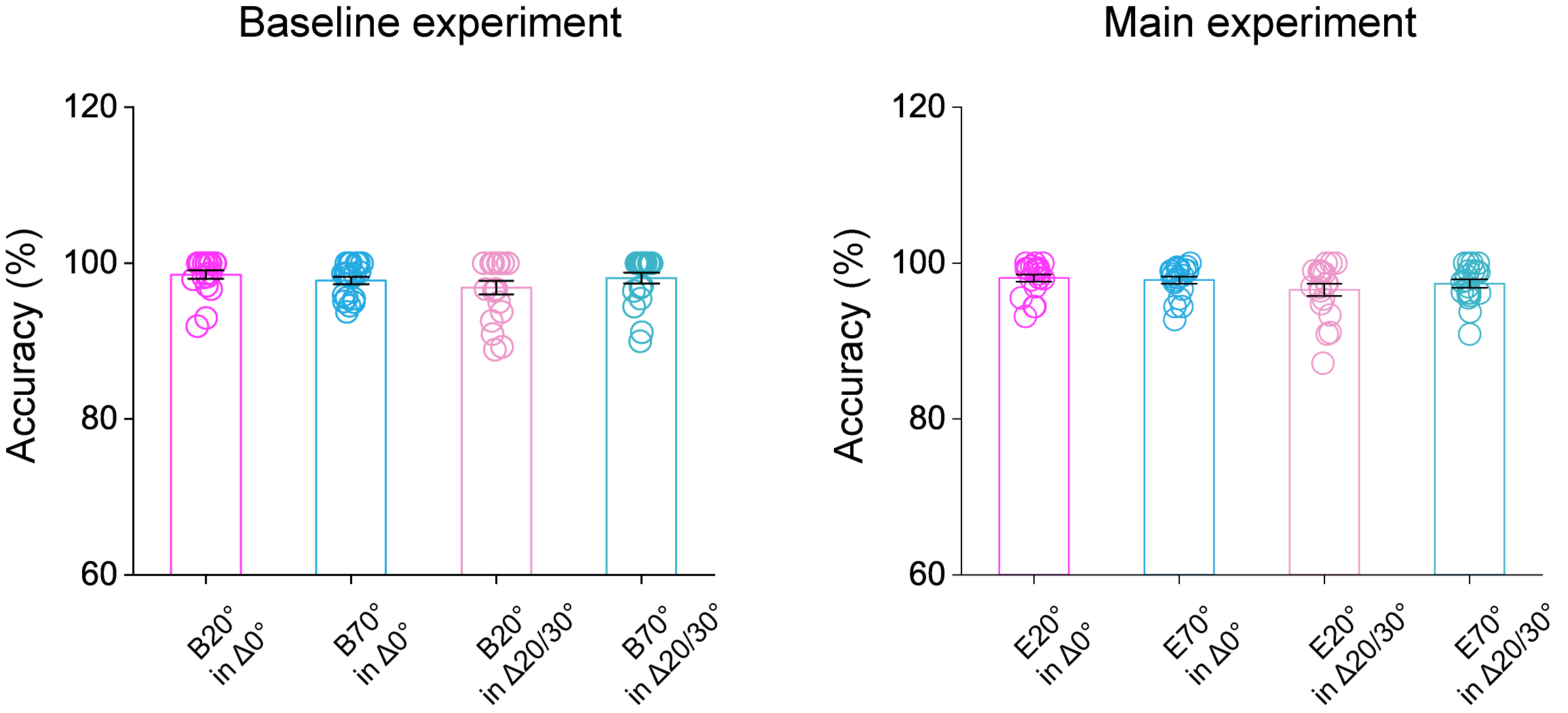

**Figure 5 – Figure supplement 1 | Accuracies of auditory tone report in orientation adjustment experiments**. Mean accuracies of auditory tone report during the baseline (left) and main (right) experiment. In both the baseline and main experiments, a repeated-measures ANOVA with the expected condition (B/E20° vs. B/E70°) and orientation distance (Δ0° vs. Δ20/30°) as within-participants factors showed that, the interaction between these two factors was not significant for either the baseline (F(1,19) = 3.330, p = 0.084, 𝜂_p_^2^ = 0.149) or main (F(1,19) = 1.509, p = 0.234, 𝜂_p_^2^ = 0.074) experiments. Error bars indicate 1 SEM calculated across participants.

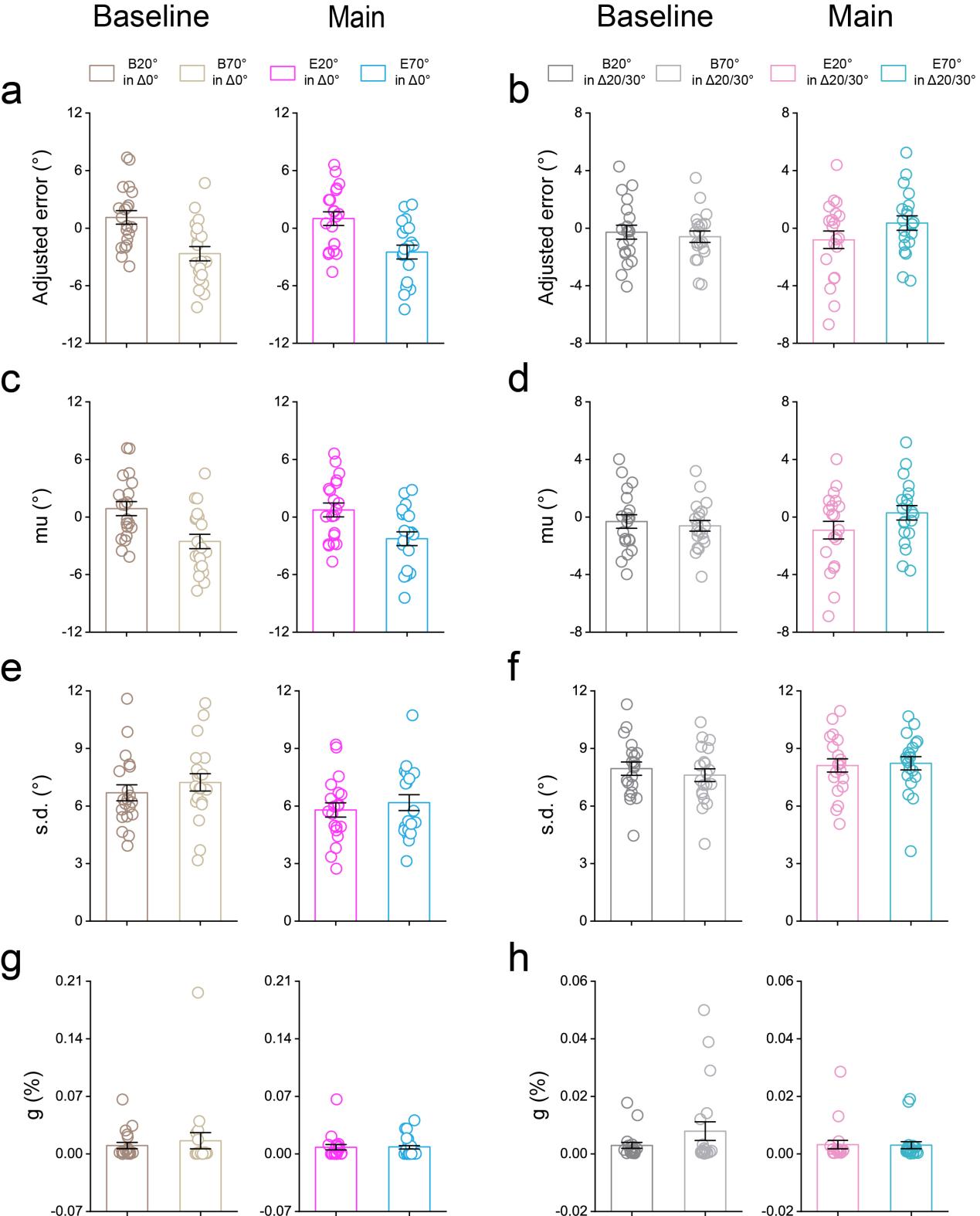

**Figure 6 – Figure supplement 1 | Adjusted errors of orientation adjustment experiments and their parameter estimates with the three-component mixture model.** The adjusted errors in both baseline (left) and main (right) experiments for expected (**a,** 20°/70°, i.e. Δ0°) and unexpected (**b,** 40°/50°, i.e. Δ20°/Δ30°) conditions. The estimated parameter *mu* in both the baseline (left) and main (right) experiments for expected (**c,** 20°/70°, i.e. Δ0°) and unexpected (**d,** 40°/50°, i.e. Δ20°/Δ30°) conditions. The estimated parameter *s.d.* in both the baseline (left) and main (right) experiments for expected (**e,** 20°/70°, i.e. Δ0°) and unexpected (**f,** 40°/50°, i.e. Δ20°/Δ30°) conditions. The estimated parameter *g* in both the baseline (left) and main (right) experiments for expected (**g,** 20°/70°, i.e. Δ0°) and unexpected (**h,** 40°/50°, i.e. Δ20°/Δ30°) conditions. Error bars indicate 1 SEM calculated across participants.

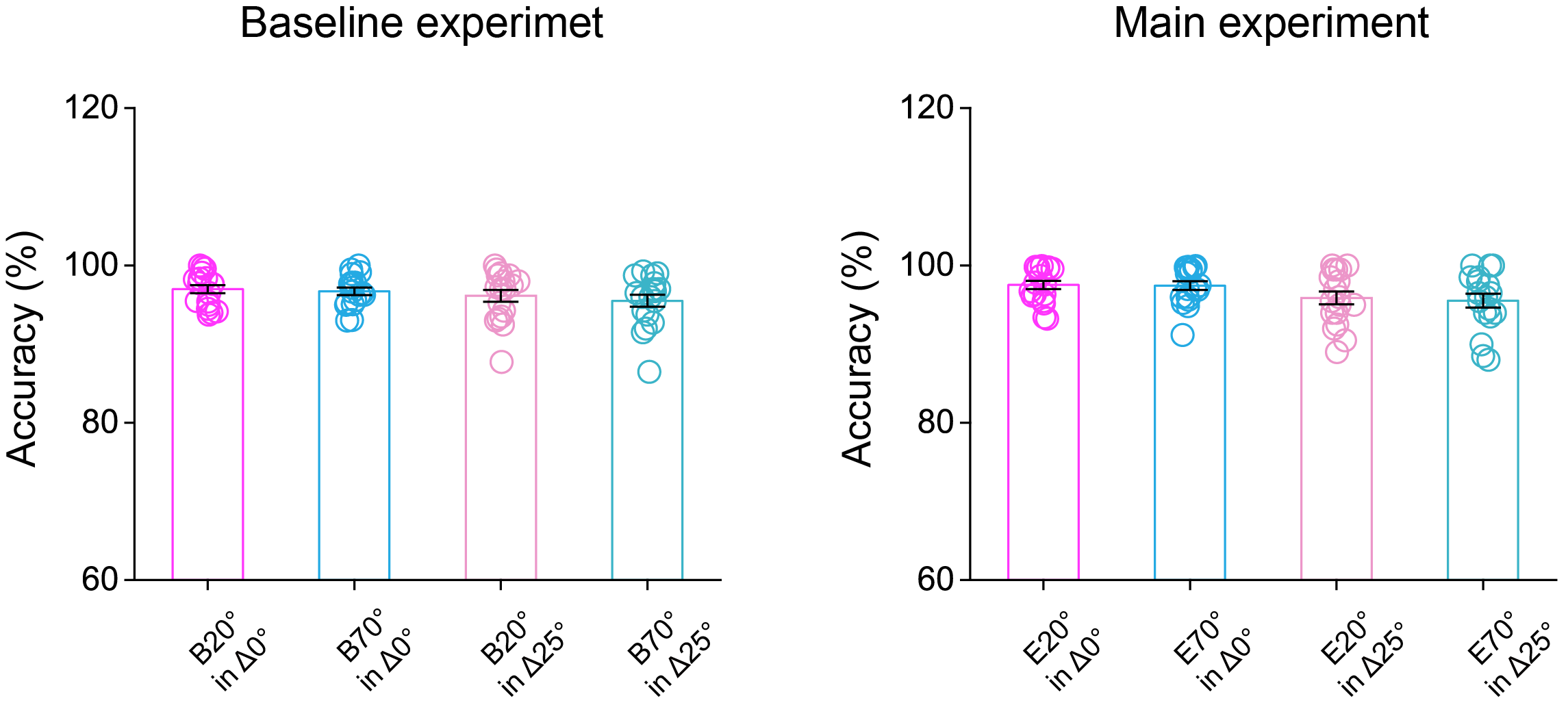

**Figure 7 – Figure supplement 1 | Accuracies of auditory tone report in orientation discrimination experiments**. Mean accuracies of auditory tone report during the baseline (left) and main (right) experiment. In both the baseline and main experiments, a repeated-measures ANOVA with the expected condition (B/E20° vs. B/E70°) and orientation distance (Δ0° vs. Δ25°) as within-participants factors showed that, the interaction between these two factors was not significant for either the baseline (F(1,17) = 0.481, p = 0.497, 𝜂_p_^2^ = 0.028) or main (F(1,17) = 0.288, p = 0.599, 𝜂_p_^2^ = 0.017) experiments. Error bars indicate 1 SEM calculated across participants.
